# Estimating extreme displacements under a Lomax generalized linear model framework: the role of sample size and threshold choice

**DOI:** 10.1101/2025.01.29.635335

**Authors:** Francisca Javiera Rudolph, José Miguel Ponciano

**Author notes:** Corresponding author: Francisca Javiera Rudolph.

## Abstract

Understanding ecological variation is fundamental to predicting the dynamics of natural systems, and it becomes increasingly more complex when rare or extreme events are involved. Extreme value statistics are a useful but underutilized tool in ecological analysis, and in this paper we showcase a simulation based study that uses extreme value distributions to estimate rare events. Focused on long-distance dispersal events, we simulate individual variation in seed dispersal events within heterogeneous populations by using finite mixture models. We employ three different approaches to model fitting extreme value distributions, and we particularly incorporate the special case of a Lomax distribution. Using a modified generalized extreme value distribution and comparing to a peak over threshold model fitting approach, we assess which models are able to capture the true underlying parameters. Additionally, we used a well-known linear bootstrap bias-corrrection and evaluate its utility to correct the bias in the estimation. Our results show that threshold models are the best at capturing the true underlying probability of long-distance dispersal models, and that the bias correction is somewhat useful, depending on the sample size. Estimating rare events in ecology is an important and statistically challenging process and in this study we provide a simple probabilistic approach to doing so.

## Introduction

Extreme value theory is a statistical discipline that focuses on describing and quantifying the stochastic behavior of a given process, unusually large or small events, beyond what has been observed, and the theory provides a class of models to extrapolate and estimate these extremes (Coles 2001). Extreme value theory has been commonly used in financial and risk analysis (Gilli and këllezi 2006), hydrology (Renard and Lang 2007, Towler et al. 2010), metereology (Burke et al. 2010), and most recently, cryptocurrencies (Gkillas and Katsiampa 2018). It is not surpriging that given the situation with the climate crisis, modeling and prediciting extreme events has become a priority (Perera et al. 2020). In the case of ecology, extreme value analysis was first suggested by Gaines and Denny (1993) given that many questions in biology were focused on analyzing the extremes rather than central tendencies. Since then, ecological approaches using this branch of statistics have been used to detect rare species (Cunningham and Lindenmayer 2005, MacKenzie et al. 2005, Mao and Colwell 2005), estimate the probability of rare events (Edwards et al. 2005), ecological distrurbances (Dixon et al. 2005, Katz et al. 2005), animal movement (Wijeyakulasuriya et al. 2019), and long-distance seed dispersal (García and Borda-de-Água 2017).

### A Brief Introduction to Statistics of Extremes in Ecology

The statistical modeling of extremes usually follows two approaches depending on the data available and the questions being asked. The first approach is a Block Maxima (BM) approach, in which the maximum (or minimum) observation for each independent study unit is selected and a Generalized Extreme Value (GEV) distribution is used to model these maxima (or minima). The second approach is focused on threshold models, Peak Over Threshold (POT), in which for all the data considered, a threshold is (arbitrarily) chosen and observations beyond that threshold are modeled using a Generalized Pareto (GP) distribution. The choice of approach is based on the characteristics of data and assumptions of independence. It is worth noting that both of these approaches are related and that given the cumulative distribution for the Generalized Pareto family with shape ξ and scale σ parameters:

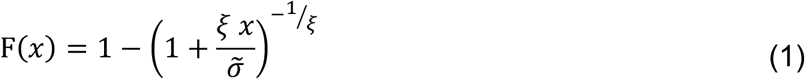

defined on {x: x > 0 and 1 + (ξ x /σ-) > 0} and with σ-= σ + ξ(x − μ) an approximation between the GEV and GP can occur, where values over a threshold excess (location, μ) modeled with a GP distribution have parameters determined by an associated GEV distribution (further details found in Coles 2001).This means, that for data arising under a BM design, if the data over a threshold excess is analyzed, the estimation of the tails under a GP model should approximate the estimation under the BM approach using a GEV distribution. This is useful in ecology, as the independence of study units and the size of the blocks is not always known, and so a POT approach to the combined data can be used, an approach previously used in genetics(Beisel et al. 2007). An example of this situation is movement data, where individual animals can belong to a social group, and thus the independence of their movements is not fully understood in relation to other group members. Additionally, a BM approach is ‘wasteful’ as it only uses the maxima or minima, discarding all other observations which might also include more information about the extremes (Coles 2001).

However, the choice of threshold in the POT approach also presents challenges, and it encompasses finding the right balance between bias and variance in the estimation, given that the higher the threshold is chosen, the less data is available for estimation and higher uncertainty in model parameters. Current approaches to selecting a threshold include visual analysis of mean excess plots or evaluating models across a large range of thresholds and selecting the highest threshold possible with the lowest variance for the estimators. These approaches have been implemented in the R package ‘extRemes’ (Gilleland and Katz 2016), but the choice of threshold is still subjective to the user and visual assessment. Newer approaches suggest the use of marginal thresholds and model evaluation (Kiriliouk et al. 2019) but have high data requirements, as they are based on stock market information, and thus ecological data does not always fulfill those requirements.

A problem that remains unexplored in the ecological literature is whether these simple probability models can accurately describe the tail behavior of complex ecological processes, and in particular, dispersal distances. Typically, the statistical features (bias, variance, mean squared error) of the estimators of a model parameter, or the tails of a distribution, are analyzed in simulation studies by generating random samples from these distributions and then assessing their bias with respect to to the true parameter values used to run these simulations. However, in ecology as in many areas in biology, these statistical models are typically misspecifications of the real and often complex biological processes, and thus whether or not we can obtain accurate descriptions of the tail behavior of complex movement or dispersal processes remains an open challenge.

In this paper, we use a computer simulation approach to assess the statistical quality of the estimated tails of a complex data-generating process by approximating them using simple extreme value distributions models. In particular, we use finite model mixtures to explicitly simulate and specify underlying heterogeneity between theoretical subpopulations. We then estimate tail probabilities using two approaches: fitting the GPD using a POT approach and fitting a special case of the GPD with only two parameters, the Lomax distribution, not using a POT approach but using all the data in the sample.

Throughout this exploration, we vary both, sample size and the right-tail quantile whose area (probability) to the right of it is to be estimated. Finally, we employ a well-known linear bootstrap bias-corrrection and evaluate its utility to correct the bias in the estimation of these right probability tails.

## Methods

### Description of Data Simulation Using a Finite Mixture Model

To describe the complexity of underlying processes generating ecological data, we used a finite mixture model framework with four mixture components and varying mixture weights. In finite mixture models (FMMs) the resulting distribution is a weighted sum of other probability densities, called components, and the weights can be determined by relevant covariates (McLachlan et al. 2019). Finite mixture models are commonly used in ecology since they allow for subpopulations to contribute to an overall probability density, or to incorporate hierarchical complexity to a resulting distribution. They have been used to describe seed dispersal kernels arising from a mix of dispersal mechanisms (Russo et al. 2006), changes in behavior and animal movement (Morales et al. 2004, Tracey et al. 2013), distribution of tree diameters (Jaworski and Podlaski 2012), and even the effect of rare species on measures of species richness (Mao and Colwell 2005). Thus, for our simulation, we chose a simple finite mixture, with four components following a Lognormal distribution and increasing weights (Figure 1):

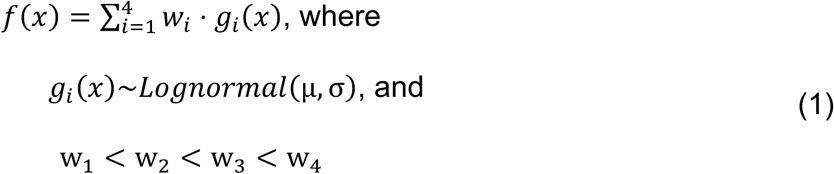

**Figure 1.**
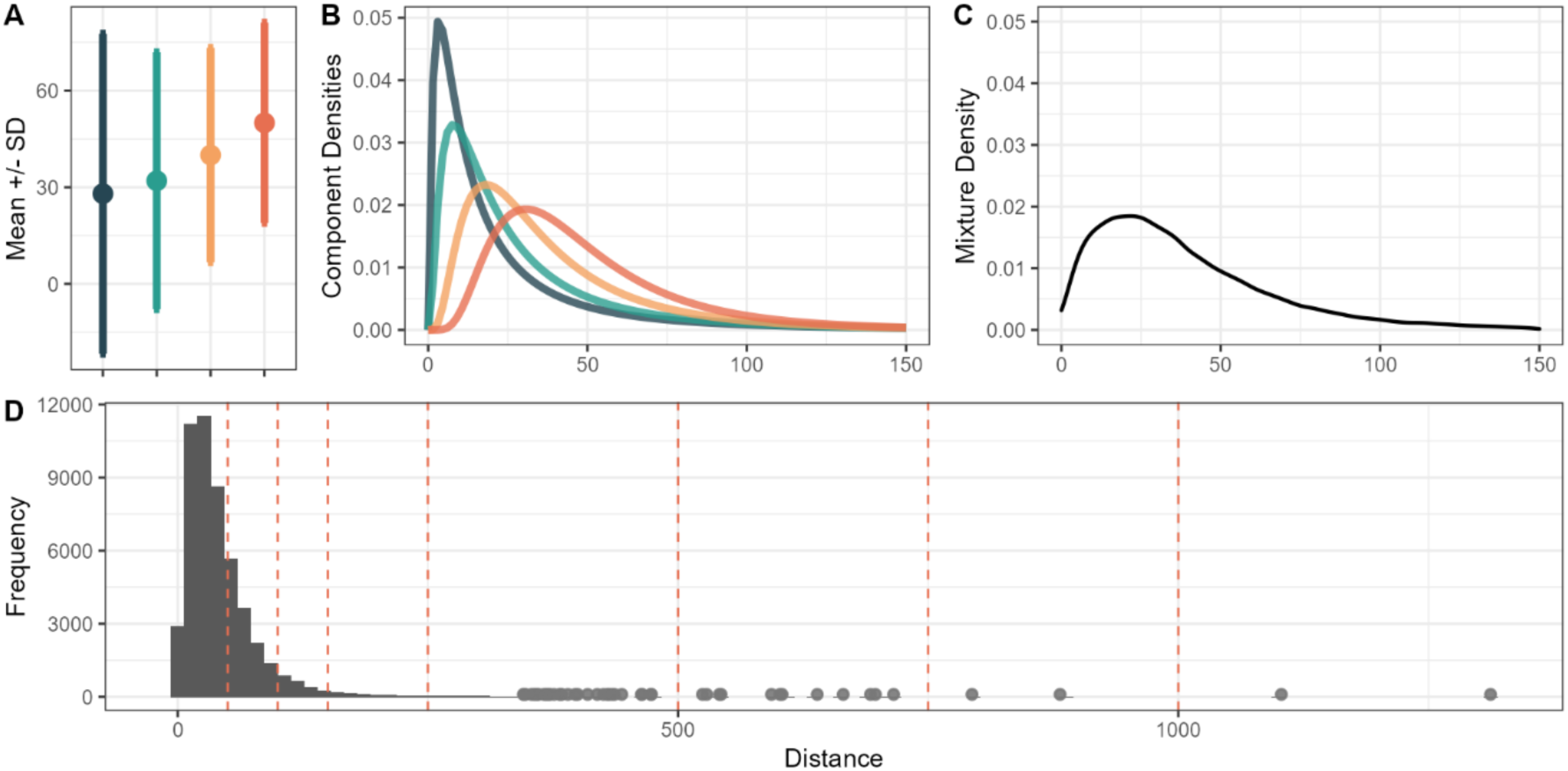
Mixture distribution scenario discussed in the main text. Panels A and B describe each of the mixture components, which are generated using a lognormal distribution function. Panel C shows the resulting mixture based on the weighted mixture components, for which weights increase from the blue component to the orange one. Panel D shows the histogram for the 50,000 samples drawn to represent the true population. Dashed red lines in panel D represent the different distance levels evaluated when estimating parameter theta, the tail of the distribution.

**Table 1.**
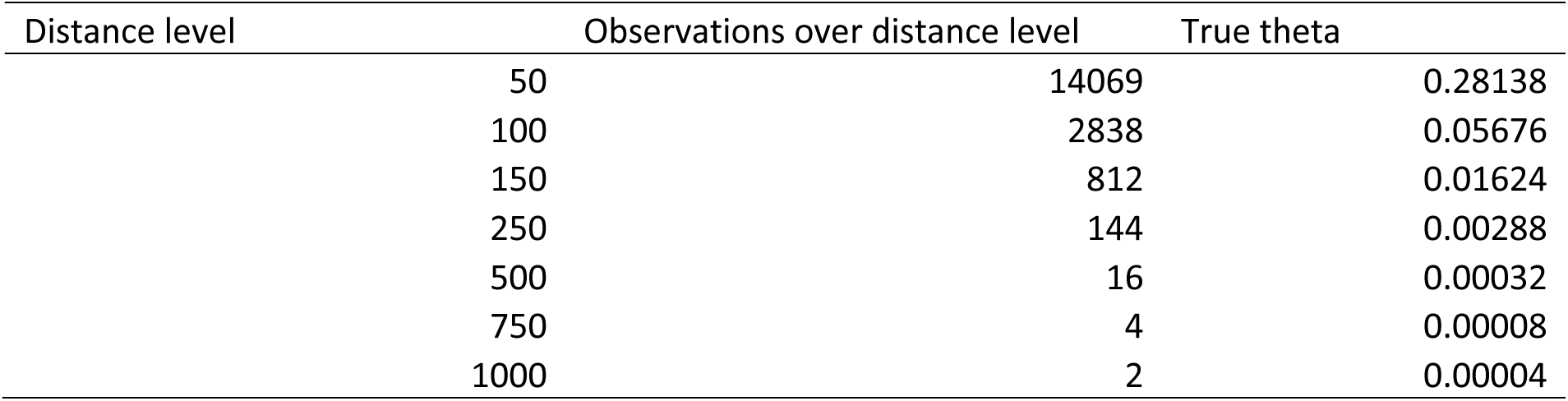
Resulting probabilities of the tail for the true population. The distance level is the distance chosen for each test. The second column shows the number of observations above the chosen distance level. The last column, theta, shows the probability of events over the chosen distance level, calculated as the second column over the total population observations, which is 50,000.

| Distance level | Observations over distance level | True theta |
| --- | --- | --- |
| 50 | 14069 | 0.28138 |
| 100 | 2838 | 0.05676 |
| 150 | 812 | 0.01624 |
| 250 | 144 | 0.00288 |
| 500 | 16 | 0.00032 |
| 750 | 4 | 0.00008 |
| 1000 | 2 | 0.00004 |

From the mixture, we generated a dataset with 50,000 distance observations, which we used as our simulated truth and will refer to as the real population from which random samples of different sizes where drawn and subsequent estimation of tail probabilities was carried.

### Testing Variation in Sample Sizes and Levels of Long-Distance Estimation

We set up our testing framework and evaluated parameter estimates of the tail for seven different levels at six different sample sizes. The underlying reasoning for this is that the estimation of rare events, whether that is species or movements, is dependent on sample size (Soetaert and Heip 1990). We assume that because the probability of extreme events is so low, that increasing sample size would increase the likelihood of extreme events in the sample and thus improve parameter estimation of heavy-tailed probability models. We established sample sizes at 80, 200, 500, 800, and 1600 samples, which were drawn from the 50,000 observations from the mixture described in the previous section. With these samples, we estimated the probability of different levels of extremes (or right-tailed quantiles) to understand how the different model fitting approaches performed. The levels tested were the same for all scenarios and with each model we calculated the probability of events happening beyond the chosen distance level. In other words, we assume *Y* is a random variable describing distances sampled, and we estimate the probability that *Y* will be greater than a chosen distance level *y*:

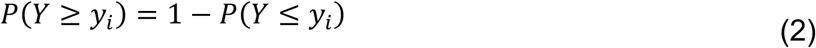

which is essentially the completementary cumulative distribution function tested at *y*_i_ for *i* = 50, 100, 150, 250, 500, 750, 1000, where *i* are the distance levels or chosen right-tailed quantiles. This function is often referred to also as survival function or reliability function, depending on the field. In the rest of the text, we refer to the true proability of *Y* being greater than the chosen distance level as theta, *P*(*Y* ≥ *y*_i_) = θ_i_, and it is calculated from the 50,000 observations generated by the mixture model in the previous section. Estimates of this probability are referred to as theta hat, θ^R^_i_, depending on the *i*^+,^ distance level being estimated. The process to estimate these tails involves fitting a probability model to the sample, estimating the parameters via maximum likelihood, and using those parameters for the survival function in order to calculate θ^R^_i_ = *P*(*Y* ≥ *y*_i_).

### Description of Probability Models Used

#### The Lomax Distribution for Individual Heterogeneity

The Lomax distribution is a heavy-tailed probability distribution, also known as Pareto type II, and is commonly used in econometrics due to its origin in modeling business failures (Lomax 1954). More recently, variations and extensions of this distribution have been applied to bladder cancer data (Rady et al. 2016) and more recently in our lab working group to model Gopher tortoise survival data. The Lomax distribution is a shifted Pareto distribution, which gives it support starting at zero, and it arises as a mixture of exponential distributions where the rate parameter is Gamma distributed. The relevance in an ecological context arises from incorporating individual variation to animal movement models. In previous work, we modeled animal movement as a random walk where step lengths were drawn from an exponential distribution. In scenarios where individual variation was incorporated, the exponential distribution’s parameter, the rate, λ, varied for each individual informed by the data. In a Lomax distribution, this rate is drawn from a Gamma distribution, where its scale, *k*, and rate parameters, α, can be used to describe this individual variability. Following this reasoning, the resulting probability density model is:

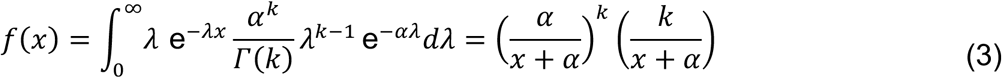

which corresponds to the probability density of a Lomax distribution. Therefore, if we set out to estimate the probability of extreme movements or displacements, while incorporating individual variability in movement, the Lomax distribution seems like a good candidate.

#### The Generalized Pareto as an Exponential-Gamma Mixture

As mentioned in previous sections, the GP distribution is a family of distributions used to describe different tail behaviors with a peak over threshold approach. The GP distribution, as described by equation 1, offers great flexibility in the description of the tail. Specifically, the value of the shape parameter ξ determines the domain of attraction under which the tail behavior falls (Beisel et al. 2007), therefore with ξ > 0, the estimated distribution falls under a Fréchet (heavy-tailed) domain, with ξ = 0, the tail behaves as an exponential distribution, and with ξ < 0, it falls under a Weibull distribution domain. The GP distribution can also arise as an Exponential-Gamma mixture, like the Lomax distribution, following:

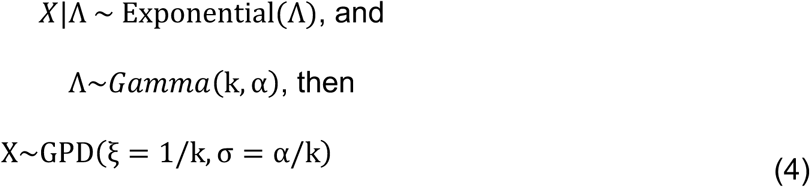

with the location parameter of the GP distribution set to zero (μ = 0), and the additional restriction that the shape parameter ξ, is positive, thus restricting the tail behavior to only two of the extreme value domains. This equivalence shows that the Lomax distribution is a special case of the GP, with support in zero, and limited to exponential and Fréchet domains for the tails. In the case of a Lomax distribution, where the GP distribution has the location parameter set to zero, the choice of threshold in the GP is circumvented.

#### Parameter estimation for Lomax and GP distributions

Parameter estimation for the Lomax and GPD models were performed using maximum likelihood with custom built functions or using the R package extRemes (Gilleland and Katz 2016). We estimated parameters for each of the samples drawn under the different sample sizes described above. Although all fitting procedures correspond to maximum likelihood, we implement three different strategies to do the model fitting.

The first strategy was based on fitting the Lomax model using the traditional likelihood function (the joing probability density function of the observations) using all the data from each sample. That is, if we had drawn a sample of 80 distances from the population data, then all 80 observations were used to fit the Lomax distribution, and each one of these distances was considered a sample regardless of individual provenance. Writing the likelihood function as the joint probability density function assumes that these distances (or observations) are independent and identically distributed random samples.

The second strategy involves phrasing maximum likelihood estimation using a more general framework that is easily adaptable to different situations. This framework corresponds to a Generalized Linear Model (GLM) fitting approach in which the distribution in question is re-parameterized so as to phrase it in terms of its mean and factors shaping its mean. Under a GLM approach, probability distributions are fitted by reparameterizing the mean of the distribution as a general function of a linear model of interest: µ = Xβ where the mean of the observations, µ, is written as a linear combination of covariates in matrix X with β coefficients. Mimicking what is typically done for Poisson-distributed GLMs, in the case of the Lomax distribution, we reparameterized the likelihood function so that its expected value, E(X) = α/(*k* − 1) is equal to Xβ.

The third and final maximum likelihood fitting strategy was implemented with the GP distribution using a peak over threshold (POT) approach, in which the choice of threshold was based on each specific sample’s quantile. Therefore, for each sample, we calculated the 0.5 quantile, and used that value as the location parameter, μ = *quantile*(0.5) to fit a Generalized Pareto distribution. As we mentioned above, threshold choice for POT models can be subjective and based on the reseracher’s visual evaluation of mean excess plots. By setting the threshold to the sample’s median, we try to evaluate how parameter estimation with an approach that takes the sample’s largest half performs when compared to the Lomax approach that considers all the data in a sample.

#### Bootstrap-Based Bias Correction for the Lomax GLM Framework

For all three fitting strategies, the estimated parameters are then used to calculate the probability of observations occurring at each of the distance levels proposed, using each of the model’s cumulative distribution functions. We compare the tail estimates of each model to the true proability of the tail as the difference in the log estimates of theta and theta hat:

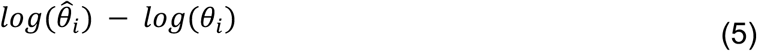

which, when considering E[*log*(θ^^^_i_)] − *log*(θ_i_), we are essentially calculating bias in the estimation of the tail. We use a bootstrap-based bias correction to improve the estimates of the tail:

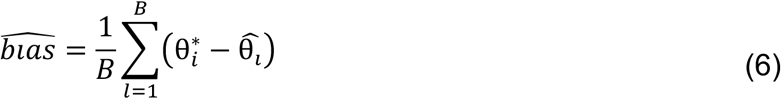

where B represents the number of bootstrap replicates of the data, and θ^*^_i_ is the tail estimate for each bootstrap sample, thus the corrected estimator, θ^-^, follows:

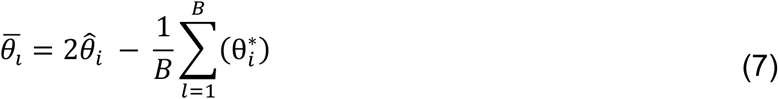

We carried out parametric and nonparametric corrections. In the nonparametric version, from the sample from which θ^^^_i_ was estimated, we sampled with replacement, refitted the Lomax model, and estimated θ^*^_i_. In the case of the parametric version, from the sample from which θ^^^_i_ was estimated, we used the alpha and k parameters estimated to generate a random sample, and from that sample, we refitted the Lomax model and estimated θ^*^_i_. Then, we compared the corrected estimator θ^-^_i_ to the true probability of the tail θ_i_.

## Results

We present in this section the results associated to the estimation of the tails using the three different fitting methods: the simple Lomax, the GLM Lomax, and the Generalized Pareto using a quantile threshold. The simulated data from the mixture distribution (Figure 4-1) was leptokurtic, with the maximum simulated distance at 1312 units, a median of 32 distance units, and a mean of 41 distance units. The true probabilities of the tail, the true θ, are shown in Table 4-1. Although the values towards the more extreme distance levels are low, the data for the population show two distances greater than 750 distances units, and two distances over 1000 distance units, out of the 50,000 simulated distances.

Tail estimates, θ^R^, were compared across the three fitting methods, and the log ratio between the estimate and the true tail value, log θ^^^/_θ_, is shown in Figure 4-2. In the case of the simple Lomax estimation, we observe that with increasing sample sizes, estimates tend to very large numbers. Further exploration of the estimates showed numerical estimation problems (Figure 4-3), where the maximum likelihood estimators of the simple Lomax model tended to infinity with increasing sample sizes, a common numerical problem when dealing with only positive parameter space. The reparameterization of the simple Lomax model into the GLM framework actually solves the numerical optimization and thus we focus on this approach instead. Panel B of Figure 4-2 shows that the tail estimates under the GLM Lomax model are biased, and such bias increases as we estimate under more extreme distance levels (150 distance units vs 1000 distance units). It also shows that increasing in sample size does not have a clear effect in correcting this bias. Finally, the Generalized Pareto POT approach shows a similar trend with regards to estimating more extreme distance levels. However, in the case of the Generalized Pareto approach, an increase in sample size does have an effect on the estimates, and reduces the variation in the estimates, although they continue to be biased.

**Figure 2.**
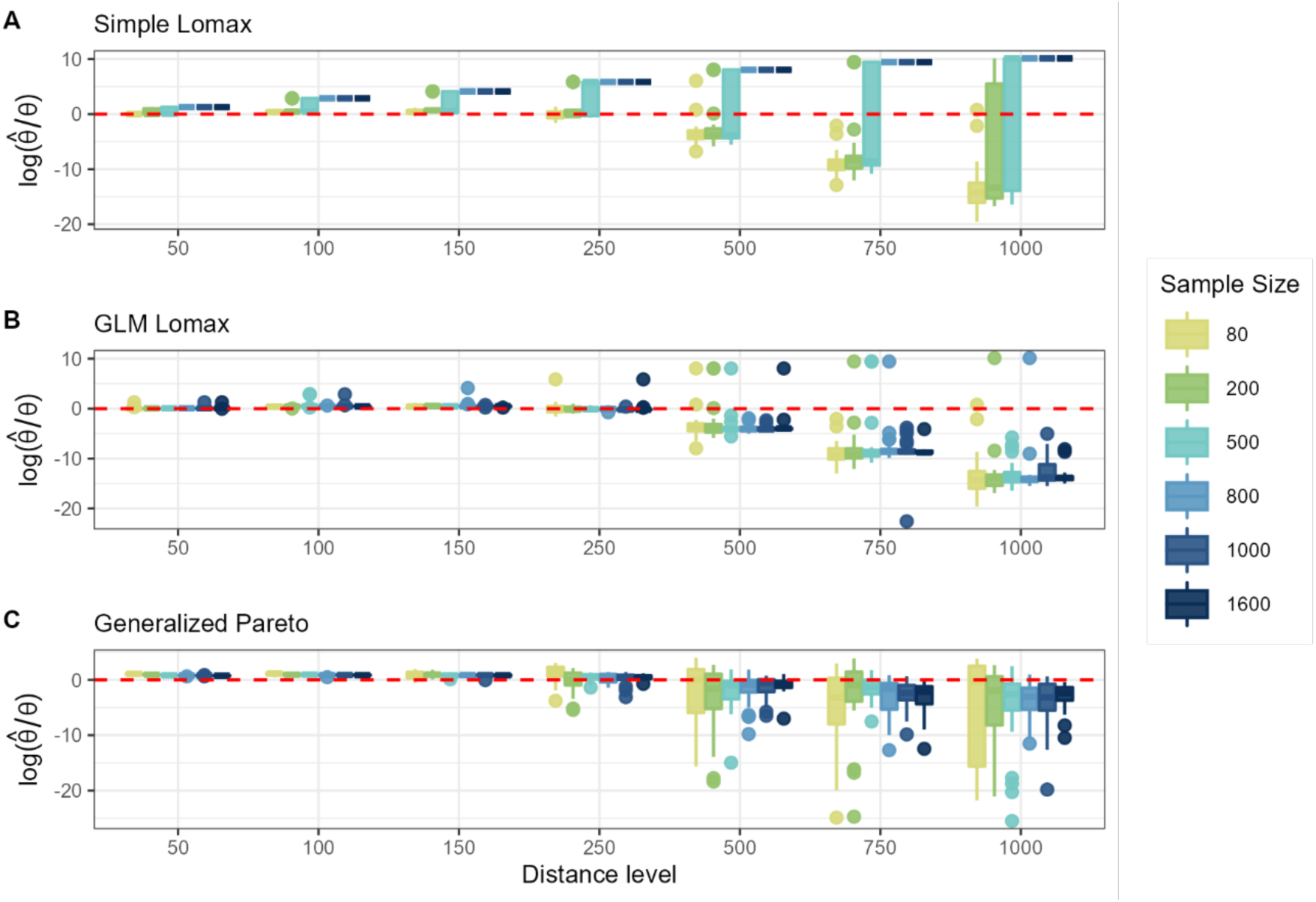
Model approximations to the true tail of the distribution. This figure shows the distribution of values of estimated thetas for each model as compared to the true theta calculated from the population. Values close to the red line, zero, represent accurate approximations, where the difference between the estimated tail and the true tail is minimal. Note the log scale on the y axis.

**Figure 3.**
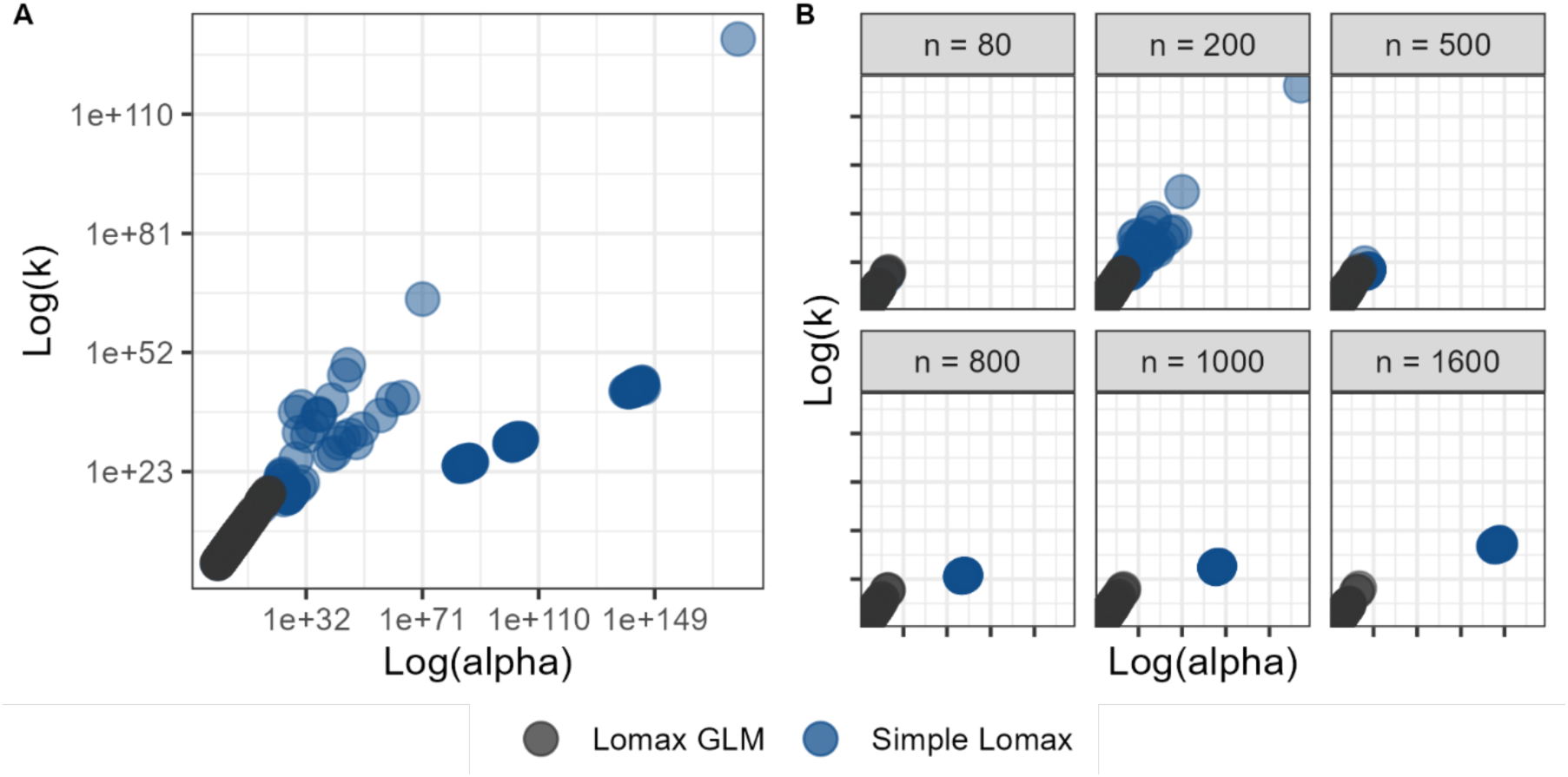
Parameter space for the simple lomax and GLM lomax. Panel A shows the aggregated parameter space occupied by both fitting approaches, the simple Lomax and GLM Lomax for all sample sizes. Note the x and y axis are in log scale and extremely high values. Panel B shows the parameter space under the same x and y axis scale but separated by sample sizes. As sample size increases, the parameter estimation for the simple Lomax tends to infinity, showing a numerical estimation problem in the process.

The bias correction performed for the Lomax distribution had no effect on the estimates, and in some cases, it produced outliers and increased the bias of the tail estimates. On the other hand, the linear bias correction for the Generalized Pareto with the quantile threshold had a beneficial effect on correcting the bias of the tail estimates (Figure 4-4). The nonparametric bootstrap correction effect varied across sample sizes and distance level estimation. At low distance estimates, both the original and corrected estimates had similar performance and were slightly biased, with overestimation of the tail. At higher and more extreme distance levels, the bootstrap correction decreased the variation in the estimators and the number of outliers. At smaller sample sizes, the bias correction tended to slightly overestimate the tail probabilities for extreme distance levels (n = 80, 200 at distance level 1000), whereas at greater sample sizes the bias correction continued to underestimate the tail probability (n = 1000, 1600 at distance level 750 and 1000).

**Figure 4.**
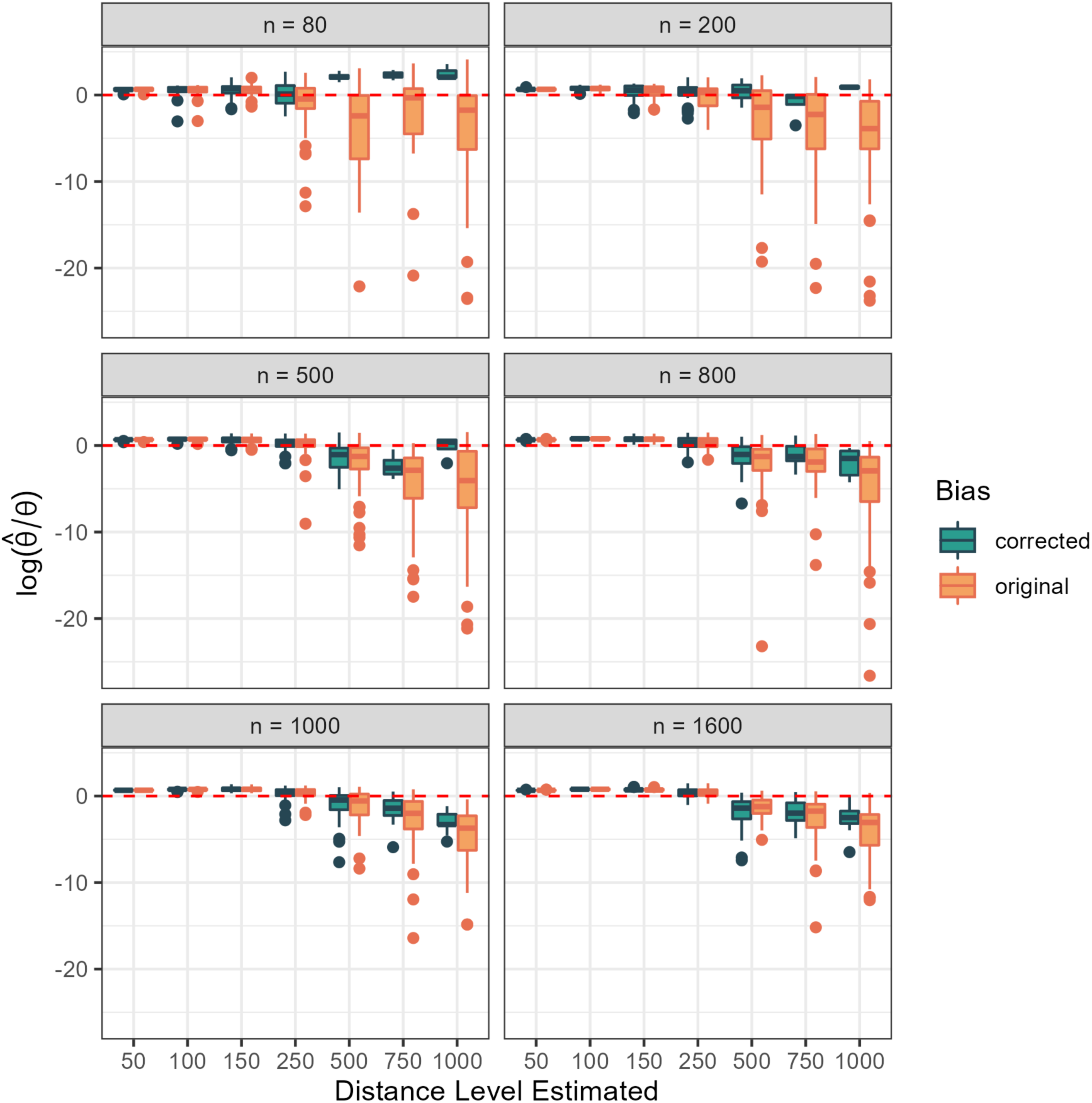
Bias corrected estimates for the Generlized Pareto Distribution. A linear bias correction using nonparametric bootstrap was performed under the GP distribution model. Each panel represents the associated sample sizes used.

## Discussion

Estimating rare events in ecology is an important and statistically challenging process. Often limited by small sample sizes, accurately estimating the probability of occurrence of rare events becomes difficult, and the smaller the probabilities of occurrence, the higher the uncertainty in these estimates. In this study, we focused on exploring different model fitting approaches to estimate the true tail of a population using probability models that constitute simple phenomenological descriptions of complex biological processes. Specifically, we evaluated how, when data is simulated from a complex mixture of behavior/distance traveling model, an increasing sample size can correct for the bias in tail estimates under a generalized linear model framework with a Lomax distribution and all the distance observations in a sample and using a Generalized Pareto distribution with a threshold set to the sample’s median, which implies using only the distance data beyond this threshold.

Overall, we found that both model fitting approaches result in biased estimators of the tails, and that bias increases when estimating the probability of the most extreme events. The simple bootstrap correction had no effect on the estimates under the GLM Lomax estimation, but the correction decreased the variation in the estimates of the GP approach. Additionally, increasing sample size did have an effect on the estimates using the GP approach. It is worth noting that both probability models, the GLM Lomax and GP, are related, and that the Lomax distribution is a special case of the GP distribution, with the threshold set to zero. This means that the Lomax distribution is essentially a simplication of the GP and by having no threshold, parameter estimation is done with a likelihood function that uses all the data in a sample. Given that both approaches result in biased estimators, we can conclude that although they are not good descriptors of the overall distribution of distances generated under the complex process simulated with the mixture distribution, they still provide reliable estimators for the tail probabilities.

Indeed, if the emphasis is put on the estimation of the tail of the distribution of the true population, one of the estimation approaches we tried, the POT for the GP model performed relatively well for various sample sizes, where the bias was mostly dependent on how far the extrapolation was performed to estimate the most extreme tail of distances. In other words, given the sampled data and the fitted model, bias occurred when trying to estimate the most extreme distances, such as the probability of having events beyond 500 distance units. This is not completely surprising given that only 22 out of the 50,000 distances were found beyond 500, and some uncertainty is expected given how rare these events are. In this case, increasing sample size had a clear effect on reducing that variation in the estimators, and in fact at a sample size of 500, the estimators had the least amount of bias (Figure 4). We observed similar trends for the other scenarios simulated (Figure 5).

**Figure 5.**
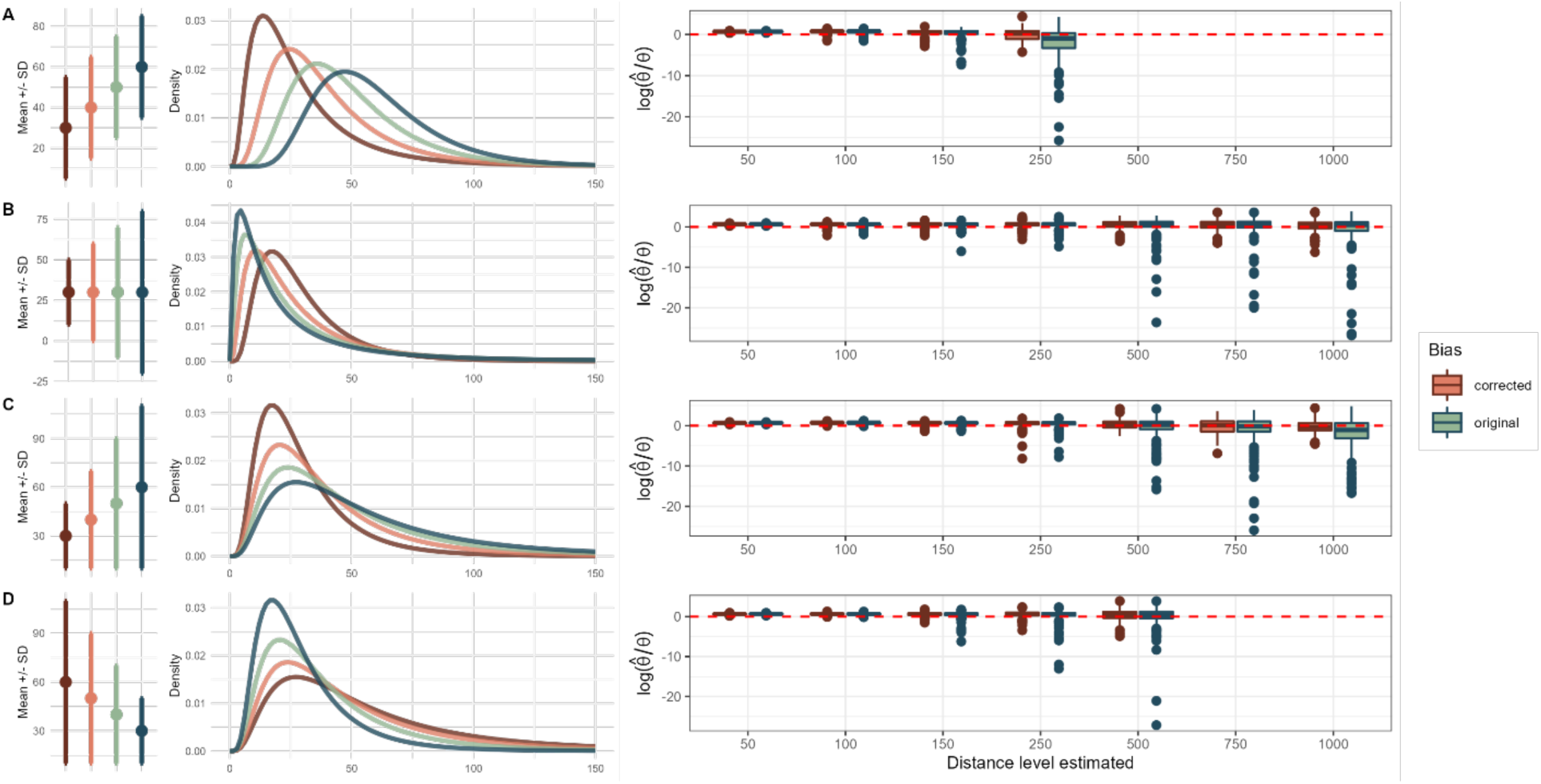
Additional mixture distribution scenarios tested for the model fitting approaches. We explored the effect of changing the mean and standard deviation of the components of the mixture. Increasing weights were assigned from left (red) to right (blue), where higher weigths contributed more to the final mixture distribution. Bias corrected estimators are shown for each of the scenarios, aggregated for all sample sizes, where bias is shown to increase when trying to estimate the true proportions of the tail with the more extreme distance levels.

Threshold models, like the Generalized Pareto presented in this work, provide useful aproximations to the tail of movement distributions by fitting models only to the observations over a high threshold. However, deciding on which part of the data to consider as the tail, in other words deciding on a threshold for the model given the data, relies on visual and exploratory techniques before fitting of models. Current approaches to selecting a threshold imply finding a balance between the bias and the variance of the estimators, done visually by fitting the models across a range of different thresholds (Coles 2001). When too low of a threshold is selected, then the model is likely to violate the assumptions for asymptotic behavior of the model, and thus leads to bias (Coles 2001). However, when the threshold selected is too high, the sample size used for the estimation is reduced and the resulting estimators show a high variance. Given the importance of the threshold selection for POT frameworks, this is an active area of research in statistics and particularly climatology (Langousis et al. 2016, Zhao et al. 2019). However, the complexity of the approaches currently being developed for threshold selection tend to be based on systems with high numbers of data and time series, such as rainfall and pollution (Tencaliec et al. 2020). In our work we suggest a simple process of threshold selection by using the median, and nonparametric bootstrap correction of the bias with the threshold set to the median of the bootstrapped samples as these settings seem to perform well. Although the simple bootstrap correction does not completely correct for the bias, it still provides close estimates of the tail probabilities, and thus, our approach can still bring novel understanding of the relevance of heavy-tailed phenomena for ecological processes when the underlying mechanisms are not known.

The mechanism-free approximation of the Generalized Pareto can provide a close enough estimate even when the complex underlying mechanisms generating the data are not completely understood. Ecology as a field is a heavily quantitative and mathematical subject (Pielou 1969), providing great opportunities for the development of biostatistical tools and modeling of species populations and communities (Gotelli and Ellison 2013). However, despite recognizing the importance of the variance in ecological processes (Benedetti-Cecchi 2003, Violle et al. 2012) the majority of models describing community and population dynamics focus on the average trends (Holyoak and Wetzel 2020). Many ecological questions are actually concerned about the extremes in a variable (Gaines and Denny 1993), and more relevant now under drastic and changing climate conditions (Perera et al. 2020), we must consider the deep consequences that extreme events, in our specific case extremes of movement and dispersal, may have on ecological dynamics. If ecologists, as a scientific community, shift a focus towards the understanding and prediction of rare events, and the impacts of these over general community dynamics, then we need to provide better guidelines regarding the estimation of these rare events and how extreme value theory is a powerful tool for this purpose. However, with the work we’ve provided in this current chapter, we are able to show that despite the complexity of the underlying generating mechanisms, using a simple and mechanism-free probabilistic model, such as the Generalized Pareto POT approach, we can gain significant understanding of the tail behavior of the distribution and closely estimate the probability of rare events.

Although understanding the complexity in biology and developing intricate models to dissect such complexity is necessary, in this work we also show the advantages of using biology-free probabilistic models to understand the occurrence of extreme events. The model fitting approaches described in this chapter are useful tools that can be readily used with available data to generate a range of projections and estimates of the tail under different scenarios, complementing other models that aim to describe the complex mechamisms giving rise to a variety of ecological dynamics.

Boxplots for each case are based on 30 runs for each combination of sample size and distance level. Panel A shows the simple lomax distribution with poor estimation with increasing sample size and distance level. Panel B shows a GLM framework of the Lomax, where poor estimation occurs towards bigger distance levels. Panel C shows the theta estimates under a Generalized Pareto distribution with a threshold set to the 0.5 quantile of the sample.

Bias corrected and original estimates are shown side by side in different colors, as a ratio of the true theta. Estimation for lower distance levels is close to zero, with some overestimation across all sample sizes. With estimation at more extreme distance levels, bias generally increases, but the correction does reduce bias, and so does increasing the sample size.

